# An explainable model using Graph-Wavelet for predicting biophysical properties of proteins and measuring mutational effects

**DOI:** 10.1101/2023.11.01.565109

**Authors:** Shreya Mishra, Neetesh Pandey, Atul Rawat, Divyanshu Srivastava, Arjun Ray, Vibhor Kumar

## Abstract

Proteins hold multispectral patterns of different kinds of physicochemical features of amino acids in their structures, which can help understand proteins’ behavior. Here, we propose a method based on the graph-wavelet transform of signals of features of amino acids in protein residue networks derived from their structures to achieve their abstract numerical representations. Such abstract representations of protein structures hand in hand with amino-acid features can be used for different purposes, such as modelling the biophysical property of proteins. Our method outperformed graph-Fourier and convolutional neural-network-based methods in predicting the biophysical properties of proteins. Even though our method does not predict deleterious mutations, it can summarize the effect of an amino acid based on its location and neighbourhood in protein-structure using graph-wavelet to estimate its influence on the biophysical property of proteins. Such an estimate of the influence of amino-acid has the potential to explain the mechanism of the effect of deleterious non-synonymous mutations. Thus, our approach can reveal patterns of distribution of amino-acid properties in the structure of the protein in the context of a biophysical property for better classification and more insightful understanding.

## I. INTRODUCTION

The vast expansion of protein sequence and structure data, combined with breakthroughs in experimental and computational methods, is advancing our understanding of links among protein structure, sequence, dynamics, and function. This knowledge is utilized to gauge how proteins interact with other molecules, peptides and potential drug targets [1]. Protein structures and functions have been investigated by a range of experimental and theoretical methods [2]-[3],[4]. Network analysis, which converts the protein structure into a network of amino-acid residue, is a computational approach that has lately been investigated. The network-based analysis of interactions [5] among amino acids in protein structure has created a unique paradigm in protein systems research [6]. Undirected networks of amino acids and their associations also known as Residue interaction networks are being used to represent protein structures [7]. According to Vendruscolo et al. [7], [8] and del Sol & O’Meara [9], residue interaction networks reveal topological information and have the potential to reveal the major biophysical properties of a protein molecule. Dokholyan et al. [10] have shown that some network features are related to protein unfolding, folding rates to some extent, and some crucial residues which act as nucleation for protein folding, tend to have substantial betweenness values in the protein transition states. Jung et. al. [11] has shown that the proportion of change in average path length normalised by protein’s size to the edge removal probability, exhibited a strong correlation with unfolding rate of proteins. Bagler & Sinha [12] discovered that the network based coefficient of assortativity shows a positive correlation with folding rates of proteins. Cusack et al. [12], [13] determined the most visited residues in networks that is dependent on shortest path and betweenness to identify important residues for protein function. Recently Graph convolutional network [14][15] has also been used for modelling protein properties.

In the evolution of organisms, the mutation in their genetic content is an essential driving force. Specific genetic mutations, e.g. single nucleotide polymorphism (SNPs), can be deleterious and cause disease in individuals. Single amino acid polymorphisms (SAPs) are due to SNPs that induce amino acid mutations and are suspected to be associated with different types of phenotypes and disorders. There is an underlying assumption that single amino acid polymorphisms linked to disease can be predicted to some extent using network topological properties. According to Li et al. [16], neighboring residues of a mutation site can help to detect its relationship with disease as such aberrations generally occur at residues with high degree or centrality. Functional modules in proteins have also been studied using network parameters. Park & Kim [17] used structure-based correlation mutation analysis to discover residues and module clusters with different functional significance in rhodopsin, encoding typical coevolutionary information in the amino acid network. However, finding the exact mechanism through which a SAP influences protein and cause disorder is not a trivial problem. Linking the property of an amino-acid residue to a biophysical property of protein can provide mechanistic insight into their mutation’s deleterious effect.

Therefore, we first developed a unique approach based on graph wavelets to engineer features that can impart tremendous improvement in modelling biophysical properties of proteins using machine learning algorithms. Further we utilized it to predict the possible effect of a SAP on a biophysical property of protein. A major advantage of graph wavelet transform is that it fractionates signal into the multi-resolution pattern of scores corresponding to detailed information of hidden modules in the graph. Our method, ProteinGW, amalgamates multispectral information of physicochemical and network-based properties of amino acids (nodes) and their interactions in the protein structure. Here, we have shown that ProteinGW derived features can be used for predicting protein folding rate (regression) and classification of transmembrane-globular, Soluble/Insoluble proteins, α-helices/β-strands. We have retrieved our data from RCSB/PDB [17][18], which represents most structures of proteins. ProteinGW also helps in measuring the effect of mutations in multispectral feature scores, which can be used to explain its deleterious effect.

There have been a few methods to predict the effect of mutations on protein function and dynamics. Most often, sequence-based methods like SIFT [19] rely on available protein sequences and their characteristics, such as, position-specific substitution matrix (PSSM). Whereas structure-based methods mostly rely on machine learning approaches which need training datasets. Capriotti et al. proposed I-mutant2.0[20], which uses features such as pH, temperature and mutation type features to train support vector machines to predict the effect of mutations. Similarly, AUTO-MUTE 2.0 [21] trains a machine-learning model using features like ordered identities of amino acids, pH, temperature and statistical contact potential in the protein structures.

Another method, DynaMut [22], has been implemented to estimate the impact of mutations on dynamics and stability of proteins due to vibrational entropy changes. Whereas a method with the name SDM [23] constructs an environment-specific amino acid substitution matrix using alignment to follow evolution rather than using machine learning to assess the effect of the mutation. Most of the previously proposed methods lack explainability about the mechanism of influence of an amino acid on the property of the whole protein. Due to lack of such clarity of the influence of an amino acid on protein property, researchers often do not trust predictive methods for new protein structures. Therefore, we further developed steps for estimating the importance of multispectral features and explainability in machine learning models in addition to predicting the effect of mutations.

## II. METHOD

### A. Weighted RIG Model

Our method builds a graph of protein molecules individually, where vertices represent the graph G(V, E) as amino acid in the protein, and edges between amino acid residues represent the distance between them. Generally, a network of residues in a protein is modelled using a residue interaction graph (RIG). In a RIG-based network, an adjacency matrix is made as per the criteria: if the physical distance between two residues (vertex) is less than or equal to a certain threshold, they are connected. We have used a weighted version of RIG-based network that can include both long and short-range interactions. Long-range interactions are important for protein function prediction since they are accountable for keeping the structure of protein unimpaired. Distance between residues is calculated using three-dimensional coordinates of atoms in protein structure available in the PDB (Protein data bank) files. We have considered the center of residue to be its alpha carbon (*Cα*) atom. Let the residues (vertices) be *V_i_* and *V_j_* with *Cα* coordinates being *(x_i_, y_i_, z_i_)* and *(x_j_, y_j_, z_j_)* in the PDB structure, the physical distance *(d_i,j_)* between them is calculated as

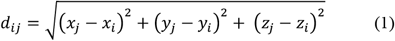

Here, i and j correspond to two residues in protein structure. These vertices are considered to be connected if this distance is less than or equal to some threshold *(r_c_)*. Hence, we have used *r_c_* = 8Å. The weighted RIG model assigns weight to the edge lying between two residue distances which is proportional to the sequence distance between the two amino-acids (residue) locations. The edge weights in the weighted RIG model are calculated as

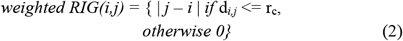

In our internal investigation, we found that the weighted RIG model tends to give better performance than the boolean RIG model. Therefore in this study we have used a weighted RIG model.

### B. Spectral Graph Theory

The weighted RIG of a protein, is just an example of the weighted undirected graph where amino acids are nodes connected by edges. A weighted undirected graph *G* comprises a defined set of vertices *V* and a positive set of weights *w: E → R* assigned to edges *E* of the graph. Let the number of vertices be defined by *N = |V|*. The weighted adjacency matrix (NxN) derived from the weighted RIG model serves as a finite weighted graph for spectral graph theory. The degree of vertices in the graph is defined as the sum of weights of edges incident on the vertex. For the degree of vertex *m*, it can also be defined as *d*(*m*) = Σ*_n_ A*_*m*,*n*_ where *A*_*m*,*n*_ is the weight of the edge between vertices *m* and *n*. With respect to the adjacency matrix. Thus, a diagonal degree matrix would be :

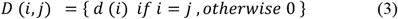

A non-normalized Laplacian operator for the RIG graph is defined by *L = D - A.* Here, we have used the normalized form of Laplacian operator given by :

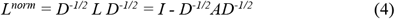

The RIG matrix is symmetric and hence can be diagonalized, which suggests eigenvalue decomposition would be *L^norm^ = UλU,* where *U* is eigenbasis matrix and *λ* is the diagonal matrix of eigenvalues of *L^norm^*. The eigenbasis matrix *U* is associated with orthonormal eigenvectors denoted as χ_l_ for 0 ≤ *l* ≤ *N* − 1. Their counterpart eigenvalues *λ_l_* satisfy *L^norm^*χ*_l_ = λ_l_* χ*_l_ .* The symmetric nature of *L^norm^* suggests that the eigenvalues are real, non-negative and can be arranged in ascending order as *0 = λ_0_ < λ_1_* ≤ *λ_2_* … ≤ *λ_N-1,_* assuming that RIG is connected.

### C. Spectral Graph Wavelet (SGWT)

The definition of SGWT [1], [24] suggests that it fixes a non-negative real-valued kernel function *g : R^+^* → *R^+,^* which is analogous to Fourier domain ψ^^*^ in the following equation

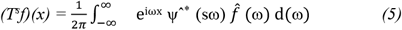

The kernel function g applied on node signal (AA features) behaves as a bandpass filter and also requires that g(0) = 0 and 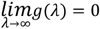. At each scale, SGWT coefficients of node signal (AA features) are produced by wavelet operators acquired as rescaled kernel function of graph Laplacian. We have used the Mexican hat filter in PyGSP (python library) for Graph signal processing applied on AA features which is defined as the second order derivative of a Gaussian.

However, for a graph Laplacian in a finite dimension, this can be achieved by eigenvectors and eigenvalues of the Laplacian matrix *(L)*. Precisely, the wavelet operator is set by *T_g_ = g(L)*. For a signal *f* (AA feature), *T_g_ f* gives wavelet coefficients at each scale (*s*). We have taken 4 scales in which AA features are decomposed into. Depending upon the action performed by this operator [24] on eigenvectors χ*_l_*, it is defined as

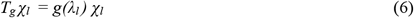

Thus, it can be said that the operator works on RIG signal *f* (AA feature) by attuning every graph Fourier coefficient as

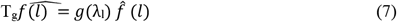

Then the wavelet operator at each scale would be 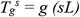. This is defined on the basis of the kernel function *(g(λ))* domain being continuous [24]. Further, the spectral graph-wavelet at scale *s*, centered on vertex *n* (AA) would be 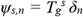 where δ_n_ is impulse on a single vertex (AA). Thus wavelet coefficients can be contemplated as inner products of *f* (AA feature) with the wavelet *ψ_s,n_* i.e

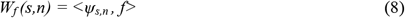

### D. Implementation Details

Our method of protein property prediction using graph wavelet (ProteinGW) first builds a weighted RIG model of each protein and puts amino acid features (physiochemical, network-based and conservation) as signals on the graph obtained (Fig. 1). Signals for the amino acid (or nodes in RIG) captured are molecular weight, hydrophobicity, amino acid frequency, conservation score, bulkiness, polarity, turn tendency, coil tendency, flexibility, partial specific volume, refractive index, compressibility, and some graph properties such as node degree, node weighted degree, clustering coefficient. Here, the node degree is the number of edges connected to the vertex and the node weighted degree is the sum of weights of edges incident on that vertex. The clustering coefficient is defined as the geometric average of subgraph edge weights i.e.,

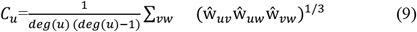

**FIGURE 1.**
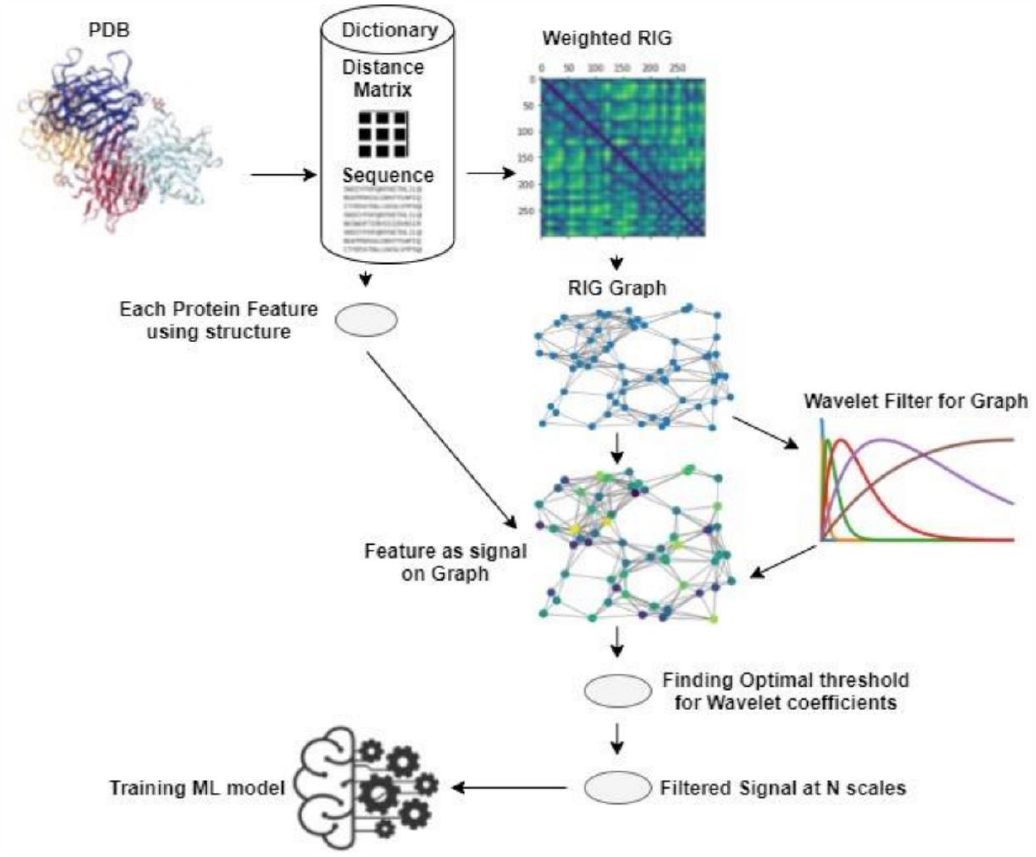
The pictorial flow chart of ProteinGW pipeline. Protein structures were collected for each protein function to be modelled. A dictionary consisting of the distance matrix, sequence, etc. was created for each protein. Weighted RIG (adjacency matrix) was created. Then, a network-specific to each protein is built using a weighted-RIG matrix. Further, physicochemical properties/network properties for each amino acid (node) are placed as a signal on the graph, the optimal threshold is calculated to threshold wavelet coefficients. Finally, a machine learning model is trained.

Here, *C_u_* becomes 0 when *deg(u) <2* and the edge weights 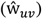 are normalized by maximum weight in the network i.e., 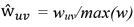.

ProteinGW yields wavelet coefficients at four scales by default. ProteinGW applies hard thresholding by finding an optimal percentile threshold on those wavelet coefficients during training of the model. This threshold is learned during the training process and is then used on the test dataset. Thus, for 15 features on 4 scales, the total feature space becomes 15*4 i.e 60. The ProteinGW fractions the dataset into training and test sets and trains machine learning models with 5-fold cross-validation. For evaluation of the predictive model, ProteinGW uses Accuracy, ROC AUC(Area under the Receiver operator Curve), Macro-F1(computed using the arithmetic mean of all the per-class F1 scores), and MCC(Matthews correlation coefficient) scores for classification tasks and R^2^(coefficient of determination) which denotes goodness of fit, RMSE(Root mean square error) for estimation of protein-folding rate. Here, we have compared random forest with other machine learning models such as XgBoost, AdaBoost, K-Nearest Neighbors, Gaussian NaiveBayes, Logistic Regression, SVM(support vector machine) for Transmembrane/Globular, all-α/all-β, Soluble/insoluble classification task. Additionally, for protein folding rate estimation using the ProteinGW approach, we compared random forest regressor with ElasticNet, Decision Tree, K-Nearest Neighbors, SVR (support vector regression), Ridge, Lasso, and Linear regressions. Further, to add explainability to the trained Random forest predictive model, ProteinGW finds feature importance at each scale for all physicochemical/network properties. Finally, change in top predictive features at disease-specific mutation sites in proteins is also studied. We have used publicly available resources [25] to determine the association of amino acid network properties with disease-associated mutation.

### E. Data Sources

For modelling folding rate, 52 single-domain two-state folding proteins were used. Information for those proteins was gathered from multiple sources [2], [26]. The same set of 52 proteins was used for alpha/beta modelling. Whereas for modeling transmembrane/globular properties, we collected 237 transmembrane and 59 globular proteins from the PDB database. For modelling solubility, we used those proteins in the list published by Han et al. [27] which have a PDB structure.

## III. RESULTS

Each amino acid has its own set of physicochemical characteristics that influence protein’s overall behavior and folding rate. As a result, extracting features based on amino acid properties is necessary and appropriate when comparing proteins and studying their function. We used known physicochemical characteristics of 20 amino acids, such as molecular weight, hydrophobicity, amino acid frequency, bulkiness, polarity, turn tendency, coil tendency, flexibility, partial specific volume, refractive index, compressibility, and some graph properties such as node degree, node weighted degree, clustering coefficient (supplementary methods) [28] and conservation score [29]. We also used the network properties of amino acids in the residue interaction graph (RIG) as features for classifying protein based on Transmembrane/Globular, soluble/insoluble, alpha/beta properties and estimating the protein folding-rate of proteins (Fig.1). ProteinGW positions signals of each property on the weighted RIG of each protein before applying graph-wavelet transform. Further, after finding a consensus cutoff to reduce noise and non-relevant components for wavelet signals, it calculates overall multispectral feature scores, which can be used in machine learning techniques. Overall, ProteinGW calculates 60 features scores where for every property of amino acid, we have 4 scores corresponding to 4 resolution levels of wavelet.

### A. The predictive power of graph-wavelet-based feature extraction

#### 1) Transmembrane-Globular Classification

First, we used our approach to predict the transmembrane-Globular property of proteins. Globular proteins have a wide range of three-dimensional structures and functions like catalysis, transport, cellular signalling etc. On the other hand, a transmembrane protein’s orientation in the membrane is always very unique. The membrane-spanning portions of the polypeptide chain that contact the hydrophobic environment of the lipid bilayer are primarily made of amino acid with nonpolar side chains which separate these domains. Because peptide bonds are polar and there is an absence of water in the bilayer, all peptide bonds are forced to establish hydrogen bonds [30]. In order to model globular/transmembrane property using graph wavelets, we collected 237 transmembrane proteins and 59 globular proteins. Multispectral features scores or, in other words, graph wavelet-based feature scores (GWFS) were calculated following the process mentioned in the methods section. Further, we used various machine learning models using GWFS and compared their performance. Different measures of evaluation of classification like accuracy, Macro F1 score, ROC-AUC (Receiver Operator Characteristic – Area Under Curve), MCC (Mathew’s correlation coefficient) are shown in Fig. 2A (supplementary table 1). Here, error bars denote standard deviation to measure variability in performance of evaluated machine learning models. These values are obtained on 5-fold cross-validation. We used a random forest-based classification model for further analysis because it has shorter error bars with an accuracy of 0.93, 0.88 Macro-F1, 0.86 ROC-AUC and 0.77 MCC scores.

**FIGURE 2.**
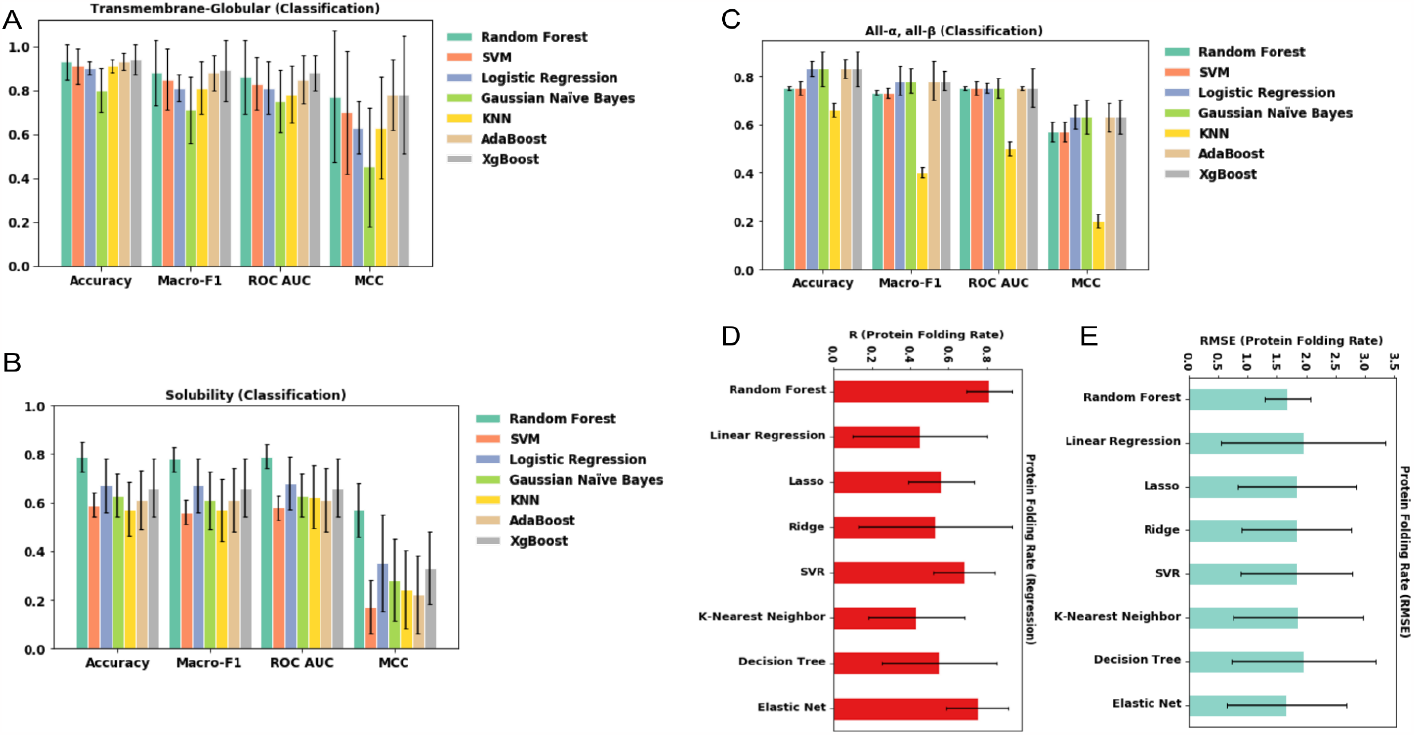
Performance of various machine learning models using 5-fold cross-validation after feature extraction using the graph-wavelet approach. (A) MCC, Accuracy, Macro-F1, ROC-AUC are compared for modeling classification of transmembrane and globular proteins. Various machine learning models i.e. XgBoost, AdaBoost, KNN, Gaussian naive Bayes, logistic regression, SVM, and random forest, are compared. (B) MCC, Accuracy, Macro-F1, ROC-AUC is compared for modelling classification of soluble and insoluble proteins by various machine learning model (C) MCC, Accuracy, Macro-F1, ROC-AUC is compared for modelling classification of all-α and all-β proteins. (D) The correlation value (R) for protein folding rate estimation is shown. Features extracted from ProteinGW are fed into the machine-learning models. ElasticNet, Decision Tree, Random Forest, KNN, SVR, Ridge, Lasso, and Linear regression are compared. (E) Root mean squared error (RMSE) for protein folding rate prediction is shown.

#### 2) Soluble-Insoluble Classification

We further tried to model the solubility of proteins using our approach. Successful recombinant protein synthesis necessitates high levels of protein expression and solubility, which is often difficult to achieve. Multiple errors during recombinant protein synthesis impede protein research, especially structural, functional, and pharmacological investigations that require soluble and concentrated protein samples [31],[32]. As a result, predicting solubility and engineering protein sequences for increased solubility is a contentious issue in research. Multiple intrinsic factors like molecular weight, amino acid composition and hydrophobicity of residues affect protein solubility [33], [34]. Therefore, we used GWFS extracted using such amino-acid properties with machine learning classifiers to predict the solubility proteins. Performance evaluation of various machine learning techniques using 5-fold cross-validation revealed that random forest was superior in classification with the accuracy of 0.79, 0.78 Macro-F1, 0.79 ROC-AUC, 0.57 MCC scores (see Fig. 2, supplementary table 2).

#### 3) All-α, all-β Classification

Another protein classification problem that we tried to solve using our approach is determining α and β content. All-α, all-β, α+β, and α/β are the four structural classifications that globular proteins fall into. All-α and all-β proteins are classified as being almost entirely made up of α-helices and β-strands. Separate segments of α-helices and β-strands (mostly antiparallel) were identified for α+β proteins composition, whereas mixed segments of α-helices and β-strands were characterized for α/β proteins (mainly parallel). We evaluated the performance of our wavelet-based feature engineering on the classification of all-α and all-β proteins. α-helices and β-strands classification is also done by deriving features in the wavelet domain with frequency cutoff using hard thresholding after finding the optimal threshold. Mostly we observed the performance of all machine learning models (features from ProteinGW are used), very close in terms of Accuracy, Macro-F1, ROC AUC, and MCC scores (supplementary table 3). However, the best performance showed 0.75 accuracy, 0.73 Macro-F1, 0.75 ROC AUC, 0.57 MCC scores. We compared the performance of our method (ProteinGW) with 3 previously published tools for predicting solubility, namely, Protein-Sol [35], GraphSol [36] and DSResSol [37](Supplementary Fig. 3). We found that ProteinGW had better performance than other 3 methods tested for solubility.

**FIGURE 3.**
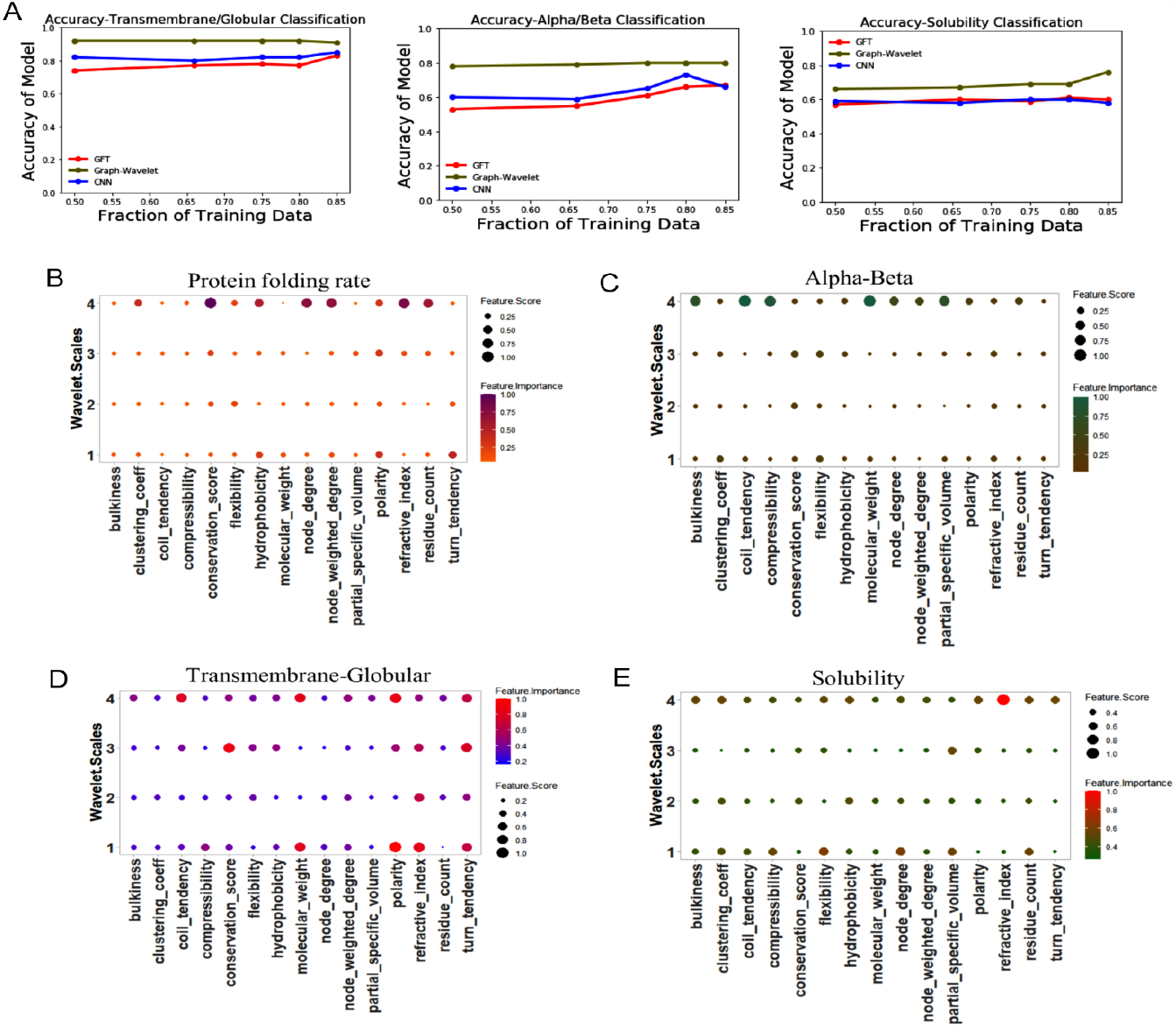
Comparison of ProteinGW with other methods and feature-weights for their amino-acid properties. (A) Accuracy of protienGW, convolutional neural network (CNN) and graph Fourier Transform (GFT) at the different number of training data points, The training fraction is reduced from 0.85 to 0.50. . The importance of the feature score was calculated using a Random forest-based model. (B) The importance of graph wavelet-based features scores (GWFS) on different scales in modelling the biophysical property of proteins. Four wavelet scales are demonstrated in estimating the protein-folding rate. The figure shows conservation score, node weighted degree, refractive index, Node degree, and residue count at scale 4 (corresponding to low frequency) are crucial, while polarity exerts more importance at scale 1. Similarly, polarity at scale 3 also appeared to have detectable importance. (C) Similarly, for transmembrane-globular, polarity, Molecular weight at scales 1 and 4, conservation score at scale 3, coil tendency at scale 4, the refractive index at scales 1 and 2 and turn tendency at scales 1 and 3 have turned out to be more important. (D) Four wavelet scales are illustrated in classifying All-α and all-β Proteins. Here, molecular weight, coil tendency, compressibility, bulkiness at scale 4 emerged to be most prominent. (E) Likewise, for the classification of soluble and insoluble proteins, the refractive index at scale 4 and flexibility, node degree, residue frequency and partial specific volume at scale 1 appeared to be more important.

#### 4) Protein Folding Rate Prediction (Regression)

After measuring the performance of our method on 3 protein-classification problems, we used proteinGW for the prediction of the fold-rate of protein. The rate of protein folding is a metric for determining how quickly (or slowly) a protein folds from its unfolded form to its native 3D structure. Protein folding rates research helps in better understanding of differences in protein folding kinetics that can contribute to illnesses like prion and Alzheimer’s disease etc [38]. We evaluated our method for modelling protein folding rate, for which we collected protein structures [12], [39] and their protein-folding rates. Their features were engineered using ProteinGW. Regression models were trained on the training dataset, and the performance was evaluated on test data with 5-fold cross-validation. RMSE and R-value of various regression models i.e., linear, lasso, ridge, SVR, KNN, random forest, decision tree, elastic net evaluated using features engineered from our method and are shown in Fig. 2D and 2E (supplementary table 4). It is evident from Fig. 2D and 2E that random forest has performed best among all models with R= 0.81 and RMSE=1.68. When we compared the RMSE for folding rate prediction reported by three other methods (Pred-PFR, FoldRate, SWFoldrate) [40], [41] (see supplementary table 5), we found that their performance was lower than proteinGW.

### B. Comparing Graph Wavelet with other graph-signal based methods

We compared our approach against other methods which can use structure-based networks and amino-acid properties on nodes to model the biophysical property of proteins. One such method is based on the graph Fourier transform, recently proposed in our report [41]. The other method we used for comparison is based on a convolutional neural network which also uses graph structure to combine signals of an input vector of quantified property of amino acids. In the field of protein function modelling, the Convolutional Neural Network (CNN) based Deep Learning has been a tremendous success, but it takes a lot of data samples to train a network of deep learning. In practice, getting a high number of training samples is too difficult, and under the constraints of a small dataset, it is most likely to overfit. We may, however, use the Fourier transform to get the amplitude of the signals required to reproduce any signal [41]. However, the Fourier transform has a basic limitation that all characteristics of a signal are global in scope. On the other hand, wavelet transform allows efficient access to localized frequency information about the signal [24]. Thus, we compared the performance of CNN, GFT (Graph Fourier Transform) (see supplementary methods), and ProteinGW when reducing the fraction of the training dataset to evaluate the robustness and consistency of these methods.

For modelling transmembrane/Globular property, we consecutively reduced the number of training samples to evaluate the performance of the model when trained with fewer data points which can be seen in Fig. 3A with Macro-F1 Score ranging from 0.87 at nearly all fractions for ProteinGW. However, the performance of graph-classifying CNN [24], [42] is comparatively less for all sizes of training data. In addition, the performance of CNN degrades progressively when this fraction is reduced slightly. We have also compared our method with the graph Fourier transform-based filtering method [41], as shown in Fig. 3A (see supplementary Fig. 1, supplementary table 6). Graph wavelets are seen to perform consistently, and performance is 92% (accuracy) even when the fraction of training data is 50%. We also performed a comparison for all-α and all-β classification and found that ProteinGW showed consistency throughout even on decreasing the number of data-points in training data (see Fig. 3A, Supplementary Fig. 1), (Supplementary Table 7). While the accuracy of ProteinGW reduced from ∼0.80 to 0.78 at 0.50 fraction of the training set, CNN model showed a larger reduction (from 0.66 to 0.59). Further, GFT accuracy ranged from 0.67 to 0.53. Further assessment was carried out for solubility. ROC AUC for ProteinGW varied from 0.73 to 0.66 as training data was reduced from 85 to 50 percent of the whole data (supplementary Fig. 1B). While for GFT, it ranged from 0.63 to 0.59 (supplementary Table 8). Similarly, 0.61 to 0.58 was the range for CNN. Overall, such analysis reveals that our method outperforms other tested methods at all sizes of the training set.

### C. Pattern and importance of graph wavelet-based feature scores of amino acids in determining protein’s biophysical property

Our approach of multispectral decomposition for analysis of residue interaction network of protein with the signal on the node has rarely been used. Hence it becomes important to study the behavior of extracted features and make insight into their pattern. We studied the trend in the importance of the scores based on the multispectral representation of physicochemical and network properties of amino-acid residues for each protein function modelling. We extracted the importance of graph wavelet-based feature score (GWFS) at 4 scales from the random forest machine learning model. Features that were most important for explaining protein-folding rate were conservation score, node-weighted degree, refractive index, node degree and residue frequency signals at scale 4 (low frequency), while hydrophobicity, and polarity have some importance at scale 1 (high frequency). The directionality of correlation of GWFS w.r.t the property of protein is shown in supplementary table 9. Similarly, conservation score and polarity at scale 3 also appeared to be mildly important (Fig. 3B). Since in wavelet transform, low frequencies (high scales) correspond to global information of a signal (or cumulation of many interactive residues with the same physiochemical property); it indicates that when a large number of residues with certain properties (like node-degree, refractive index) are connected with each other, then the folding rate is high. Whereas, if there are smaller groups of tightly connected residues with such property or when the structure is composed of smaller modules, the folding rate is slow.

We also calculated the importance of amino acid properties and their distribution pattern on the residue graph for the classification for transmembrane and globular proteins (see Fig. 3C). Properties that appeared to be most salient based on random forest-based classifiers were polarity and molecular weight at scales 1 and 4. Such a result can be explained by the fact that proteins with higher globularity have polar amino acids at the surface with less interaction with each other (higher value of high-resolution signal, scale-1). In contrast, transmembrane proteins have nonpolar amino acids dispersed on the surface such that polar amino-acid interact with each other at the core, leading to a higher value of low-frequency spectrum of polarity (scale-4) of polarity (see Fig. 3C and supplementary table 9). A similar argument could be given for coiled-coil tendency, whose low-frequency component seems to be important. It indicates that there could be larger modules of interacting residues with a high coiled-coil tendency in globular proteins. Other interesting patterns in the importance of GWFS for modelling transmembrane/globular properties were for conservation score at scale 3, the refractive index at wavelet scales 1 and 2 and turn tendency at scales 1 and 3.

Similarly, for the classification task of alpha and beta proteins (shown in Fig. 3D), molecular weight, coil tendency, compressibility, and bulkiness at scale 4 emerged to be foremost in their classification. It is well known that coiled-coil structures stabilize alpha-helices, so larger modules of interacting residue with high coil tendency would be more common in alpha proteins. However, for differentiating soluble and insoluble proteins, our model showed refractive index at scale 4 (low resolution) appeared to be more important than other feature scores. Other features which showed detectable importance were flexibility, node degree, residue frequency, and Partial specific volume at scale 1 (see Fig. 3E). Overall, the importance of GWFS corresponding to a few residue properties and wavelet scales provides insights into how different types of amino acids interact to make different modules.

### D. Graph wavelet-based features add explainability to the effect of mutations on protein’s biophysical property

Our approach provides an advantage of measuring the multispectral intensity of the score of predictive and important features at every amino acid. Such as, for the classification of transmembrane/globular protein, we can get the graph-based multispectral score of the most predictive feature (polarity) on every amino acid. Such information of feature score from every amino acid can be used to explain the importance of every amino acid in determining the class of protein as a transmembrane or globular. Therefore, we further investigated the utility of our methods for finding the possible mechanism of the effect of disease-associated mutations. We curated the disease-associated mutations in proteins used in our study for modelling globularity and protein folding rate with the help of Uniprot [1] and HuVarBase (HUmanVARiantdataBASE) databases [25] (supplementary table 10). Out of the mutations mentioned in supplementary table 10, we could find an explanation of their possible effect using our model, as elaborated below.

#### 1) Case studies based on Transmembrane/Globular properties of Proteins

We first investigated the percentile of GWFS of each amino acid for top properties identified using feature-importance provided by random forest classifiers. Further, we investigated if high GWFS (high percentile) for top predictive features for an amino acid for globularity prediction could explain their mechanism of influence. Such as, one of the globular proteins Lamin A 17-70 coil1A dimer stabilized by C-terminal capping (PDB ID:6YF5) is mentioned to have a mutation in arginine to Glycine/Proline (at Position:25), which causes disease Emery-Dreifuss muscular dystrophy 2 [25] (autosomal dominant version) (EDMD2). Our initial analysis shows that the associated specific location (at position: 25) has a moderately high polarity-based average (mean of all layers) GWFS (∼ 75th percentile) among all amino acids (supplementary table 11). We also found another mutation in residue 50 in the PDB structure Lamin A (6YF5), which is associated with EDMD2(autosomal dominant). The residue 50 in LAMIN A structure (PDB id: 6YF5) also has moderately high polarity-based average GWFS (∼ 75 percentile) (see Fig. 4A). When we estimated layer-wise GWFS for polarity, both the residue locations (25 and 50) in 6YF5 had a rank below 72 percentiles in the fourth layer of wavelet (Supplementary Table 12), which is important for modelling globularity (see Fig. 3C and Supplementary Table 9).

**FIGURE 4.**
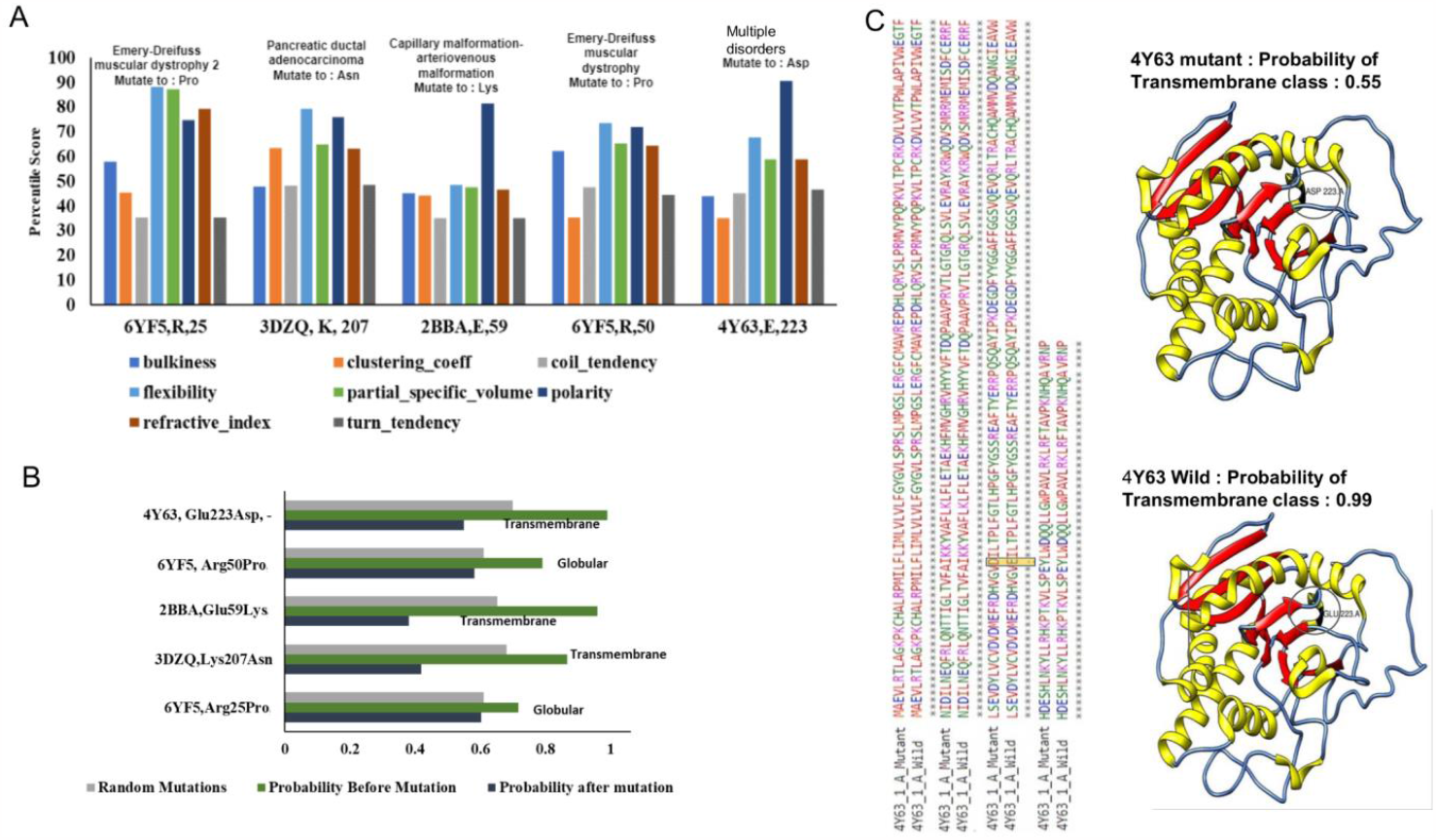
Relevance of transmembrane/globular property in the effect of disease-causing mutation (A) Percentile Scores of top features i.e. polarity, bulkiness, flexibility, refractive index, clustering coefficient, partial specific volume, turn tendency, and coil tendency for proteins structures with PDB id : 3DZQ (EPHA3, Lys207Asn*, Pancreatic ductal carcinoma*), 2BBA (EPHB4, Glu59Lys*, Capillary malformation-arteriovenous malformation*), 6YF5 (LMNA, *Arg50Pro, Emery-Dreifuss muscular dystrophy*), 6YF5 (LMNA, Arg25Pro*, Emery-Dreifuss muscular dystrophy*) and 4Y63 (ABO gene, Glu223Asp). For every feature, it represents the average value across all four levels of the wavelet spectrum. (B) Probability scores of proteins (same as in subpart a) before and after mutation of the original class of the proteins. For 2BBA and 3DZQ it shows a probability of being transmembrane. For comparison with the null model, the average change in probability for mutations at 10 random sites is also shown for every protein. (C) Visualization of 4Y63 mutation site and probability of belonging to transmembrane class before and after mutation.

We also found high polarity-based GWFS at a residue in Human EphA3 kinase protein, which is most frequently mutated in lung cancer and other cancers, including melanoma, glioblastoma, and pancreatic [43], [44] and hepatocellular carcinoma and head and neck squamous cell carcinoma [44]. In the structure of Human EphA3 kinase domain in complex with inhibitor AWL-II-38.3 (PDB ID: 3DZQ) the mutation in residue position 207 from Leucine to Asparagine (Leu207asn) is known to be associated with the disease Pancreatic ductal adenocarcinoma (Position:207). The polarity-based GWFS at residue 207 is high among all amino-acids (85.63). Among our set of PDB structures used to model globularity, we also found another protein EphB4 kinase domain inhibitor complex (PDB ID: 2BBA), with a mutation (Glutamic acid to Lysine at position 59) known to be associated with Capillary malformation-arteriovenous malformation. Our result shows a high percentile score of polarity (81.422) of residue position 59 in the structure of ephB4 kinase. The percentile scores of major top predictive features for the above-discussed proteins are shown in Fig. 4A. The average percentile and wavelet level-wise rank of polarity-based GWFS is shown in Supplementary Table 11 and Supplementary Table 12.

Further, we investigated the impact of mutations through a change in predicted globularity using our trained model for the transmembrane/globular property of proteins. The estimated probability for Lamin A structure (6YF5) being globular decreased only by 15 percent (from 0.71 to 0.60) on replacing arginine by proline (at Position:25) (Fig. 4B). Whereas for Lamin A structure (PDB ID: 6YF5) the estimated probability for being globular decreased from 0.79 to 0.58. We also simulated mutations on 10 randomly chosen residue locations on Lamin A structure (6YF5) and estimated changes in the predicted probability for globularity. The average change in the probability of globularity by 10 random mutations was almost equal to replacements at residue positions 25 and 50. On the other hand, for transmembrane protein structure, Histo-blood Group ABO System Transferase (PDB Id:4Y63) the estimated probability after mutation at 223 from Glu to Asp was reduced by 40 percent (from 0.99 to 0.55). At the same time, the drop in the probability of transmembrane property due to 10 random mutations in Histo-blood Group ABO System Transferase (PDB Id:4Y63) was ∼25 percent. Similarly, for transmembrane proteins structure with PDB IDs 3DZQ(EphA3), 2BBA(EphB4), the probability of being trans-membrane reduced from 0.86, 0.96 to 0.41, 0.38 on mutations at residues 207 and 59, respectively. The substantial reduction of more than 50 percent in the probability of being transmembrane due to mutation Lys207Asm in EphA3 provides support for a possible loss of surface localization of EphA3 reported to be influencing cancer development [44]. The simulated mutations at 10 random sites in both protein structures 3DZQ and 3BBA caused a reduction of only 25 percent. We have also shown a visualization of mutation in Histo-blood Group ABO System Transferase (4Y63, Glu223Asp) and the probability of the protein belonging to the transmembrane class before and after mutation. Overall, our result highlighted the loss of transmembrane property due to mutation at site with higher polarity-based GWFS as a possible cause of the deleterious effect of corresponding mutations (Glu223Asp in Histo-blood Group ABO System Transferase, Lys207Asm EphA3 and Glu59Lys in EphB4). We further confirmed our model with experimentally validated effects of mutations in protein EPHA3 (pdb:3DZQ) published by Liasbeth et al., [44]. Our predictions for the effect of mutations on the transmembrane property of protein EPHA3 were concordant with experimentally measured changes in cell-surface localisation (see Supplementary Fig. 4).

#### 2) Case Studies on the effect of mutations on Folding rate

We also curated known disease-associated mutations in proteins which we used for modelling folding rate. Here we investigated percentile scores for important features in Protein folding rate prediction after taking the average of their values at four levels of the wavelet spectrum. One such protein is tumor suppressor protein P16INK4A (PDB id: 2A5E) with a mutation at residue 53 from methionine to isoleucine (Met53Ile) linked to Melanoma, cutaneous malignant. The corresponding residue position 53 has a high percentile score for conservation score (76.76) and a relatively higher percentile score for flexibility (see Fig. 5A) (supplementary table 12 and supplementary Table 13). We also investigated other mutations in protein P16INK4A, namely glucine to glycine at residue location 119 (Glu119Gln) reported in biliary tract tumors and valine to alanine at 95 (Val95Ala), causing Non-small cell lung carcinoma. We also considered the case of the crystal structure of Bovine Carbonic Anhydrase II (CAII) (PDB id: 1V9E) as the mutation in its human homologue at location 94 (histidine to tyrosine) is linked to Osteopetrosis. The corresponding location in Bovine CAII protein has slightly high GWFS based on conservation, the refractive index, which appeared as an important feature for folding rate (see Fig. 5A).

**FIGURE 5.**
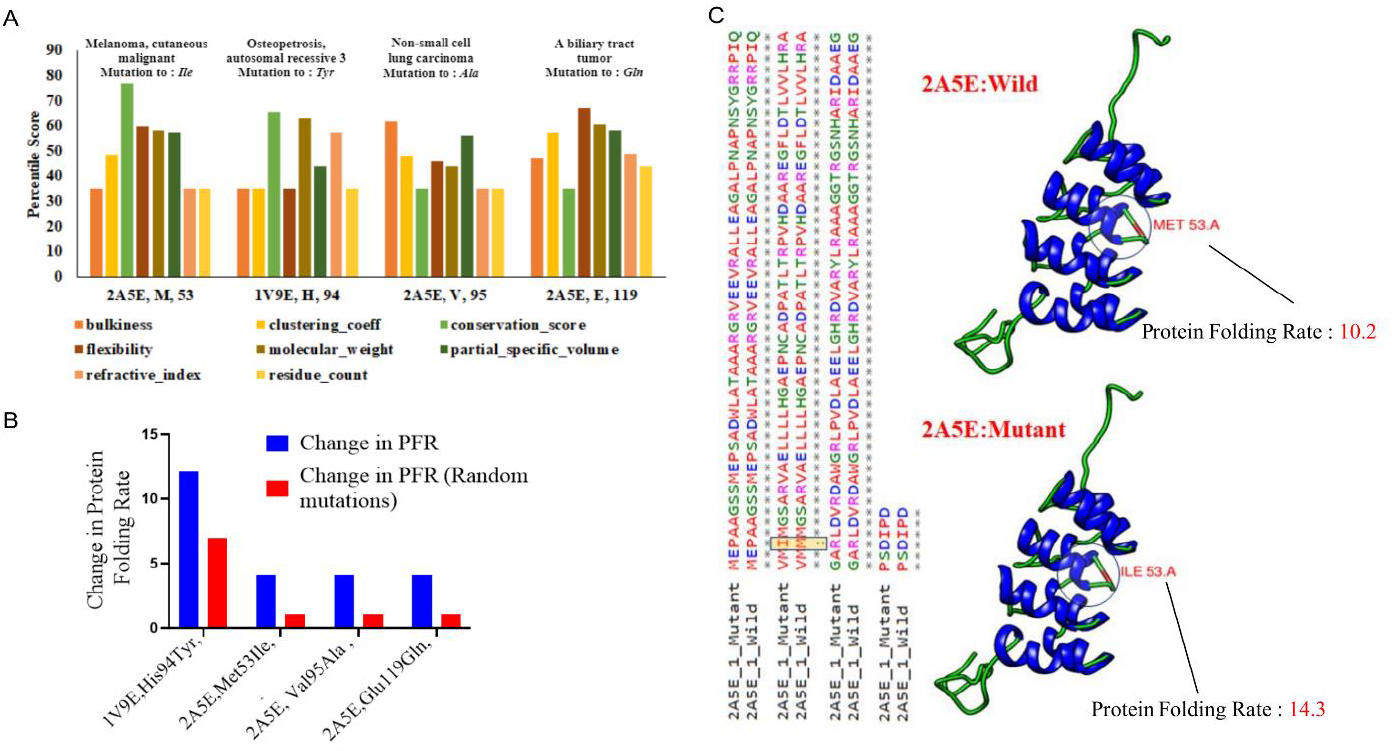
Finding relevance of folding rate in the effect of disease-causing mutation (A) Percentile Scores of top features i.e. bulkiness, conservation score, flexibility, refractive index, clustering coefficient, partial specific volume, residue frequency, molecular weight for proteins 2A5E(Met53Ile*, Melanoma, cutaneous malignant*), 1V9E(His94Tyr*, Osteopetrosis, autosomal recessive 3*), 2A5E(Val95Ala*, Non-small cell lung carcinoma*), and 2A5E(Glu119Gln*, A biliary tract tumor*). (B) Change in Protein folding rate of proteins (same as in subpart a) before and after mutation. The average change in probability for mutations at 10 random sites is also shown for every protein. (C) Visualization of 2A5E mutation site (Met53Ile*, Melanoma, cutaneous malignant*) and protein folding rate predicted, actual before mutation, and protein folding rate after mutation.

We further used our pre-trained model to check whether there is a change in the estimated folding rate of protein upon mutation at mentioned sites leading to disease. The changes in Protein folding rate due to simulated mutations are shown in Fig. 5B. The change for 2A5E(Met53Ile) was recorded to be 4.10809, and for 1V9E(His94Tyr) it was 12.11 (526% change). Similarly, for 2A5E(Val95Ala) and 2A5E(Glu119Gln) the change is 4.10527 and 4.10739 (Supplementary Table 14). The effect on folding rate by mutations at random sites in both protein structures 2A5E and 1E9E was not comparable to deleterious mutation studies here. Thus, our method generated the hypothesis of the impact on folding rate by the deleterious mutations investigated here, which can help researchers in designing relevant experiments.

## IV. DISCUSSION

Our study demonstrates a generalisable use of spectral graph-wavelets in extracting features of proteins. Here we have not targeted to study only one particular property of protein but focused more on our novel framework using which relevant features can be extracted from protein structures. ProteinGW amalgamates the amino-acid properties and residue network topology to extract features useful for modelling various physical properties of proteins. Spectral graph wavelet helps ProteinGW impart the understanding of the global and local importance of a feature for better prediction for a specific property of the proteins. However, our method also finds an optimal threshold for each residue wavelet coefficient for the refinement of noise in the signals. We have shown the performance of our method for Transmembrane/Globular, Soluble/Insoluble, all-α/all-β Proteins classification and estimation of protein-folding rate. Nevertheless, the proposed method can be used for modelling any protein property. Our method has also outperformed CNN and feature extraction using Graph Fourier transform to predict four protein properties. To evaluate the folding rate prediction, we also compared RMSE with three other methods.

One major drawback of using the graph Fourier-based approach is that it does not allow estimating the effect of each amino acid on the biophysical property of proteins. On the other hand, graph-wavelet-based method allows the investigation of the effect of each residue property leading to predicting its association with known deleterious effects (disease-causing mutation). Thus, ProteinGW exhibits explainability to predictive modelling using wavelet features to impart a better understanding of feature importance for each property individually. If the training-set is available, ProteinGW has the potential to highlight the importance of amino-acids w.r.t given the property of protein. The result of our analysis suggests possible links between proteins EPHA3 (PDB id: 3DZQ), LMNA (PDB id: 6YF5), EPHB4 (PDB id: 2BBA) and polarity score at residue sites 207, 50 and 59, respectively which could be responsible for their known associations with different disorders. Especially for cell membrane proteins EPHB4 and EPHA3, the effect of the simulated mutations at a single relevant residue location (Glu59Lys and Glu207Asm) were much higher. Such results show how a perturbation in the signal of physicochemical properties of a single amino acid without changing the residue-interaction graph can have a wide effect. The underlying cause of such a wide effect could be the 3-dimensional modular arrangement of physicochemical effects such as hydrophobicity and the strategic location of residues linking different modules. Notice here a module describes the community of connected residues sharing the same physicochemical property signals. Hence, besides the structure of the residue-interaction graph, the pattern of connected components of the physicochemical signal at different resolutions also decides the importance of a residue. Our method can capture such patterns in the structure of proteins to generate a hypothesis related to the effect of exonic mutations that could be further used for a detailed study.

Our approach based on graph wavelet transform of residue interaction graph also opens an avenue to classify proteins. Structure-based classification of proteins (SCOPE) has remained a relevant problem, at the same time, finding the cause of the influence of mutations is also important for the current post-genome-sequencing era. ProteinGW holds promise for resolving both the problems with its elegant use of graph-wavelet. Finally, graph-wavelet based feature-extraction from protein structures using ProteinGW could also be helpful in the unsupervised classification of proteins. Therefore, proteinGW could prove to be a useful resource for many researchers. A limitation of proteinGW is that it is dependent on a training set for reporting the possible effect of mutations. Such as to estimate the possible effect of a mutation on solubility, we need a training-set which includes the structure of the soluble and insoluble protein. However, given the current availability of high-confidence predictions of protein structures (such as alpha fold [44], [45], our method could be highly useful for predicting multiple kinds of the physical properties of proteins. Thus, with the support of structure prediction methods like “alpha fold” our approach could help in resolving the issue of estimating the mechanism of the effect of exonic mutations genome-wide. Hence in future, we hope to collect more training-set for multiple other types of protein properties to achieve a better overview of the effect of mutations.

## REFERENCES

[1] UniProt Consortium, “UniProt: a worldwide hub of protein knowledge,” Nucleic Acids Res., vol. 47, no. D1, pp. D506–D515, Jan. 2019.

[2] E. Alm and D. Baker, “Matching theory and experiment in protein folding,” Curr. Opin. Struct. Biol., vol. 9, no. 2, pp. 189–196, Apr. 1999.

[3] B. K. Shoichet, W. A. Baase, R. Kuroki, and B. W. Matthews, “A relationship between protein stability and protein function,” Proc. Natl. Acad. Sci. U. S. A., vol. 92, no. 2, pp. 452–456, Jan. 1995.

[4] A. Rives et al., “Biological structure and function emerge from scaling unsupervised learning to 250 million protein sequences,” Proc. Natl. Acad. Sci. U. S. A., vol. 118, no. 15, Apr. 2021, doi: 10.1073/pnas.2016239118.

[5] G. Hu, W. Yan, J. Zhou, and B. Shen, “Residue interaction network analysis of Dronpa and a DNA clamp,” J. Theor. Biol., vol. 348, pp. 55–64, May 2014.

[6] J. W. Heal, G. J. Bartlett, C. W. Wood, A. R. Thomson, and D. N. Woolfson, “Applying graph theory to protein structures: an Atlas of coiled coils,” Bioinformatics, vol. 34, no. 19, pp. 3316–3323, Oct. 2018.

[7] J. Zhou, W. Yan, G. Hu, and B. Shen, “Amino acid network for the discrimination of native protein structures from decoys,” Curr. Protein Pept. Sci., vol. 15, no. 6, pp. 522–528, 2014.

[8] M. Vendruscolo, N. V. Dokholyan, E. Paci, and M. Karplus, “Small-world view of the amino acids that play a key role in protein folding,” Phys. Rev. E Stat. Nonlin. Soft Matter Phys., vol. 65, no. 6 Pt 1, p. 061910, Jun. 2002.

[9] A. del Sol and P. O’Meara, “Small-world network approach to identify key residues in protein-protein interaction,” Proteins, vol. 58, no. 3, pp. 672–682, Feb. 2005.

[10] N. V. Dokholyan, L. Li, F. Ding, and E. I. Shakhnovich, “Topological determinants of protein folding,” Proc. Natl. Acad. Sci. U. S. A., vol. 99, no. 13, pp. 8637–8641, Jun. 2002.

[11] J. Jung, J. Lee, and H.-T. Moon, “Topological determinants of protein unfolding rates,” Proteins, vol. 58, no. 2, pp. 389–395, Feb. 2005.

[12] G. Bagler and S. Sinha, “Assortative mixing in Protein Contact Networks and protein folding kinetics,” Bioinformatics, vol. 23, no. 14, pp. 1760–1767, Jul. 2007.

[13] M. P. Cusack, B. Thibert, D. E. Bredesen, and G. Del Rio, “Efficient identification of critical residues based only on protein structure by network analysis,” PLoS One, vol. 2, no. 5, p. e421, May 2007.

[14] V. Gligorijević et al., “Structure-based protein function prediction using graph convolutional networks,” Nat. Commun., vol. 12, no. 1, p. 3168, May 2021.

[15] X.-M. Zhang, L. Liang, L. Liu, and M.-J. Tang, “Graph Neural Networks and Their Current Applications in Bioinformatics,” Front. Genet., vol. 12, p. 690049, Jul. 2021.

[16] Y. Li et al., “Predicting disease-associated substitution of a single amino acid by analyzing residue interactions,” BMC Bioinformatics, vol. 12, p. 14, Jan. 2011.

[17] K. Park and D. Kim, “Structure-based rebuilding of coevolutionary information reveals functional modules in rhodopsin structure,” Biochim. Biophys. Acta, vol. 1824, no. 12, pp. 1484–1489, Dec. 2012.

[18] H. M. Berman et al., “The Protein Data Bank,” Nucleic Acids Res., vol. 28, no. 1, pp. 235–242, Jan. 2000.

[19] P. C. Ng and S. Henikoff, “SIFT: Predicting amino acid changes that affect protein function,” Nucleic Acids Res., vol. 31, no. 13, pp. 3812–3814, Jul. 2003.

[20] E. Capriotti, P. Fariselli, and R. Casadio, “I-Mutant2.0: predicting stability changes upon mutation from the protein sequence or structure,” Nucleic Acids Res., vol. 33, no. Web Server issue, p. W306, Jul. 2005.

[21] M. Masso and I. I. Vaisman, “AUTO-MUTE 2.0: A Portable Framework with Enhanced Capabilities for Predicting Protein Functional Consequences upon Mutation,” Adv. Bioinformatics, vol. 2014, p. 278385, Aug. 2014.

[22] C. H. Rodrigues, D. E. Pires, and D. B. Ascher, “DynaMut: predicting the impact of mutations on protein conformation, flexibility and stability,” Nucleic Acids Res., vol. 46, no. W1, pp. W350–W355, Jul. 2018.

[23] A. P. Pandurangan, B. Ochoa-Montaño, D. B. Ascher, and T. L. Blundell, “SDM: a server for predicting effects of mutations on protein stability,” Nucleic Acids Res., vol. 45, no. W1, pp. W229–W235, Jul. 2017.

[24] D. K. Hammond, P. Vandergheynst, and R. Gribonval, “Wavelets on graphs via spectral graph theory,” Appl. Comput. Harmon. Anal., vol. 30, no. 2, pp. 129–150, Mar. 2011.

[25] K. Ganesan, A. Kulandaisamy, S. Binny Priya, and M. M. Gromiha, “HuVarBase: A human variant database with comprehensive information at gene and protein levels,” PLoS One, vol. 14, no. 1, p. e0210475, Jan. 2019.

[26] L. Yu, Y. Zhang, I. Gutman, Y. Shi, and M. Dehmer, “Erratum: Protein Sequence Comparison Based on Physicochemical Properties and the Position-Feature Energy Matrix,” Sci. Rep., vol. 7, p. 46787, May 2017.

[27] X. Han, X. Wang, and K. Zhou, “Develop machine learning-based regression predictive models for engineering protein solubility,” Bioinformatics, vol. 35, no. 22, pp. 4640–4646, Nov. 2019.

[28] S. Kawashima, P. Pokarowski, M. Pokarowska, A. Kolinski, T. Katayama, and M. Kanehisa, “AAindex: amino acid index database, progress report 2008,” Nucleic Acids Res., vol. 36, no. Database issue, pp. D202–5, Jan. 2008.

[29] J. A. Capra and M. Singh, “Predicting functionally important residues from sequence conservation,” Bioinformatics, vol. 23, no. 15, pp. 1875–1882, Aug. 2007.

[30] B. Alberts, Molecular Biology of the Cell. 2004.

[31] R. M. Kramer, V. R. Shende, N. Motl, C. N. Pace, and J. M. Scholtz, “Toward a molecular understanding of protein solubility: increased negative surface charge correlates with increased solubility,” Biophys. J., vol. 102, no. 8, pp. 1907–1915, Apr. 2012.

[32] Q. Hou, R. Bourgeas, F. Pucci, and M. Rooman, “Computational analysis of the amino acid interactions that promote or decrease protein solubility,” Sci. Rep., vol. 8, no. 1, p. 14661, Oct. 2018.

[33] D. D. Lawson and J. D. Ingham, “Estimation of solubility parameters from refractive index data,” Nature, vol. 223, no. 5206, pp. 614–615, Aug. 1969.

[34] B. K. Bhandari, P. P. Gardner, and C. S. Lim, “Solubility-Weighted Index: fast and accurate prediction of protein solubility,” Bioinformatics, vol. 36, no. 18, pp. 4691–4698, Sep. 2020.

[35] M. Hebditch, M. A. Carballo-Amador, S. Charonis, R. Curtis, and J. Warwicker, “Protein-Sol: a web tool for predicting protein solubility from sequence,” Bioinformatics, vol. 33, no. 19, pp. 3098–3100, Oct. 2017.

[36] J. Chen, S. Zheng, H. Zhao, and Y. Yang, “Structure-aware protein solubility prediction from sequence through graph convolutional network and predicted contact map,” J. Cheminform., vol. 13, no. 1, p. 7, Feb. 2021.

[37] M. Madani, K. Lin, and A. Tarakanova, “DSResSol: A Sequence-Based Solubility Predictor Created with Dilated Squeeze Excitation Residual Networks,” Int. J. Mol. Sci., vol. 22, no. 24, Dec. 2021, doi: 10.3390/ijms222413555.

[38] G. M. Ashraf et al., “Protein misfolding and aggregation in Alzheimer’s disease and type 2 diabetes mellitus,” CNS Neurol. Disord. Drug Targets, vol. 13, no. 7, pp. 1280–1293, 2014.

[39] M. M. Gromiha, A. M. Thangakani, and S. Selvaraj, “FOLD-RATE: prediction of protein folding rates from amino acid sequence,” Nucleic Acids Res., vol. 34, no. Web Server issue, pp. W70–4, Jul. 2006.

[40] C. C. H. Chang, B. T. Tey, J. Song, and R. N. Ramanan, “Towards more accurate prediction of protein folding rates: a review of the existing Web-based bioinformatics approaches,” Brief. Bioinform., vol. 16, no. 2, pp. 314–324, Mar. 2015.

[41] D. Srivastava, G. Bagler, and V. Kumar, “Graph Signal Processing on protein residue networks helps in studying its biophysical properties,” Physica A, vol. 615, no. 128603, p. 128603, Apr. 2023.

[42] A. J.-P. Tixier, G. Nikolentzos, P. Meladianos, and M. Vazirgiannis, “Graph classification with 2D convolutional neural networks,” in Artificial Neural Networks and Machine Learning – ICANN 2019: Workshop and Special Sessions, in Lecture notes in computer science. Cham: Springer International Publishing, 2019, pp. 578–593.

[43] V. Corbo et al., “Mutational profiling of kinases in human tumours of pancreatic origin identifies candidate cancer genes in ductal and ampulla of vater carcinomas,” PLoS One, vol. 5, no. 9, p. e12653, Sep. 2010.

[44] E. M. Lisabeth, C. Fernandez, and E. B. Pasquale, “Cancer somatic mutations disrupt functions of the EphA3 receptor tyrosine kinase through multiple mechanisms,” Biochemistry, vol. 51, no. 7, pp. 1464–1475, Feb. 2012.

[45] J. Jumper et al., “Highly accurate protein structure prediction with AlphaFold,” Nature, vol. 596, no. 7873, pp. 583–589, Aug. 2021.

